# Nuclear position modulates long-range chromatin interactions

**DOI:** 10.1101/2022.05.31.494201

**Authors:** Elizabeth H. Finn, Tom Misteli

## Abstract

The human genome is non-randomly organized within the cell nucleus. Spatial mapping of genome folding by biochemical methods and imaging has revealed extensive variation in locus interaction frequencies between cells in a population and between homologs within an individual cell. Commonly used mapping approaches typically examine either the relative position of genomic sites to each other or the position of individual loci relative to nuclear landmarks. Whether the frequency of specific chromatin-chromatin interactions is affected by where in the nuclear space a locus is located is unknown. Here, we have simultaneously mapped at the single cell level the interaction frequencies and radial position of more than a hundred locus pairs using high-throughput imaging to ask whether the location within the nucleus affects interactions frequency. We find strong enrichment of many interactions at specific radial positions. Position-dependency of interactions was cell-type specific, correlated with local chromatin type, and cell-type-specific enriched associations were marked by increased variability, sometimes without a significant decrease in mean spatial distance. These observations demonstrate that genome organization relative to itself and relative to nuclear landmarks are closely interwoven.

**Significance Statement:** A gene’s nuclear environment is defined by its distance to other genes as well as its distance to nuclear structures such as the nuclear periphery. While both of these features have been shown to be important for gene function, they are often studied separately. We performed the first systematic analysis comparing these two features. We determined that at the level of single chromosomes they are correlated, suggesting that genome organization relative to itself and relative to nuclear landmarks are closely interwoven.

## Introduction

The mammalian genome within the cell’s nucleus is non-randomly organized (1). The spatial organization of the genome is guided by two general principles. The first is chromatin folding, which refers to the position of a genomic locus relative to other loci mediated by the interaction of the chromatin fiber over short and long genomic distances. The second is nuclear organization, which refers to the 3D position of a genomic locus relative to nuclear bodies, for example the nuclear lamina or nuclear speckles (1–4).

Chromatin folding is largely thought to arise via the combined effect of two molecular processes: loop extrusion and heterochromatin self-association (5, 6). Sequence-specific chromatin loops are generated by loop extrusion, in which the cohesin motor protein complex spools the chromatin fiber through its ring until it meets the boundary protein CTCF, and the cohesin complex stalls and stabilizes associations between two distal CTCF sites in the form of a loop (7–10). Repetition of this process in the region between two convergent CTCF sites ultimately creates domains of contiguous chromatin marked by an enrichment for interactions between loci within the domain as compared to loci in neighboring domains, termed topologically associating domains (TADs) (11). Cohesin-dependent loop extrusion is counterbalanced by homotypic interactions within chromatin that drive the separation of the genome into euchromatic and heterochromatic compartments, referred to as A and B, via self-association (5, 6). It is believed that this tendency for chromatin to self-organize is driven by phase-separation of chromatin-associated proteins, especially within heterochromatin (12–15). Genome folding changes upon differentiation (16, 17), and specific loops are disrupted in disease (18), suggesting a functional role for genome architecture.

Higher order nuclear organization, in contrast, is created by the non-random location of genomic loci within the 3D space of the cell nucleus. A convenient indicator of 3D genome location is the radial position of a locus, denoting the degree of proximity to the nuclear lamina (19–21). The nuclear lamina is a broadly repressive region marked by the presence of constitutive heterochromatic domains referred to as lamin-associated-domains (LADs) (22, 23). LADs are uniquely marked by characteristic chromatin modifications, in particular histone H3K9 dimethylation (24), which is likely recognized by proteins found at the lamina to target regions to peripheral positions. However, the specific proteins recognizing the mark are unknown, and radial positioning of different loci appears to require different factors (25). Radial position in numerous examples correlates with gene activation, as many genes tend to move towards the nuclear interior from the periphery as they are transcriptionally activated, for instance immunoglobulins (26, 27), Hox genes (28), and globin genes (29). Furthermore, repositioning to the nuclear periphery often, but not always, silences genes (30–32), and radial position of specific genes is disrupted in cancer (33, 34), suggesting a role for nuclear organization in gene regulation and cell fate.

It thus appears that both genome folding and nuclear organization have ramifications for genome function. Both genome folding and nuclear organization are also marked by high cell-to-cell variability (35). Single-cell high-throughput chromosome conformation capture (Hi-C) and imaging-based studies have shown that pairwise interactions between any two genome loci generally only occur at low frequencies, and genome folding is marked by exceptional plasticity (36–39). Furthermore, it is likely that even the most conserved and specific features, such as TADs, are dynamic structures representing the average of many different conformations (40–42). Radial position similarly occurs in a probabilistic manner, such that any locus can be found at any position when examined in individual cells, and it is only in populations of cells that preferences arise (3, 43). Furthermore, LADs interact frequently with the lamina, by definition, but also the edge of the nucleolus and can sometimes be found in the nucleoplasm itself (4). Thus, while the principles of genome folding and nuclear organization are generally conserved, they are also both marked by extensive variability at the level of cells and even homologs.

Several models suggest that overall genome organization is informed by a combination of both genome folding and nuclear organization. Computational models suggest that self-association of heterochromatin, and its association with proteins at the nuclear lamina, are both required and sufficient to drive the separation of A and B compartments and their relative position within the nucleus (44). Furthermore, integrative models of genome architecture involving both Hi-C (i.e. genome folding) and lamina-DamID (i.e. nuclear organization) data yield significant additional insights compared to models based on Hi-C or lamina-DamID data alone (45). However, as these computational models depend largely on separate datasets or measurements, the relationship between chromatin folding and nuclear organization within a single nucleus is as yet unclear.

To address whether 3D nuclear position affects the frequency of a given chromatin interaction, we undertook a systematic analysis of both radial position and pairwise spatial distances at more than 100 locus pairs in human foreskin fibroblasts (HFF) and 25 locus pairs in human embryonic stem cells. We discovered that many chromatin interactions show preferred radial positions, and that loss of preferential positioning within the nucleus coincides with loss of enriched pairwise interaction frequencies. Relative enrichments in interaction frequency in a cell type could be caused by either a decrease in mean distance or an increase in variability of distance. Our observations suggest that nuclear position plays a role in promoting chromatin interactions and in this way contributes to large scale genome architecture.

## Results

### Determination of radial position of chromatin-chromatin interactions

We set out to ask whether the likelihood of intrachromosomal chromatin interactions to occur is related to their radial position within the nucleus. To this end, we selected a set of 137 probe pairs among a total of 69 loci on human chromosomes 1, 4, 17, and 18 and located in the A as well as the B compartment (Fig 1A, Table S1). Loci were selected to maximize the range of Hi-C capture frequencies at a given genomic distance, as described previously (37). All probe pairs tested were between loci on the same chromosome and no inter-chromosomal interactions were analyzed due to their very low frequency (11). Genomic distances between loci in tested interaction pairs ranged from ~52 kb to 235 Mb (Fig 1A, Table S2). As a control for the accuracy of distance measurements, a single locus was stained using three separate fluorophores, to experimentally determine the practical resolution of the microscope used. 95% of signals were within 187 nm, and the median distance was 91 nm with a standard deviation of 465 nm (Fig S1A). With the exception of the closest-spaced probe pair which was separated by 52 kbp center-to-center, all interaction pairs showed higher separation distances. For example, probes separated by 224 kbp had a median spatial distance of 180 nm and a standard deviation of 257 nm, statistically significantly different from our control pair (t-test: 6*10^−4^) (Fig S1A). This behavior confirmed that the resolution of the microscope and probes was sufficient to detect spatial separation between the locus pairs tested, as previously described (37). For each interacting pair, we mapped the radial and Euclidean position of both interaction partners by DNA-FISH using BAC probes (see Materials and Methods). For the diploid HFF and H1 embryonic stem cells used in this study, we considered only data from cells with exactly two segmented spots for each interaction partner, in wells where at least 60% of cells had exactly two segmented spots for each interaction partner, and we analyzed at least 250 cells per interaction pair. This dataset included a total of 932,261 analyzed spot pairs with a median of 3,333 spot pairs per probe pair (Table S2).

**Figure 1:**
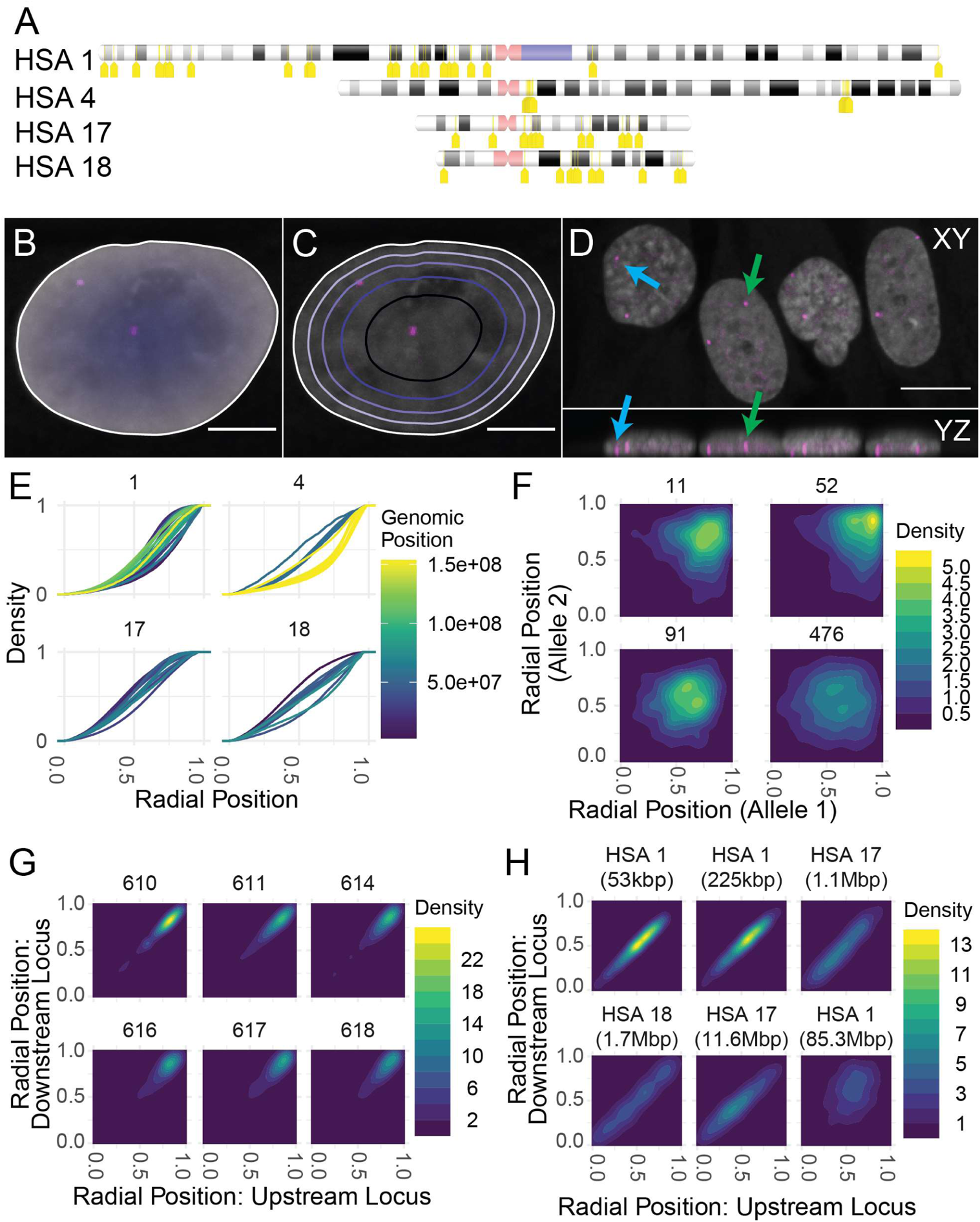
Determination of radial position of genome regions. A: Schematic location of genome probes used. Probes are indicted in yellow; centromeres in pink, pericentromeric regions in purple. B: Diagram of normalized Euclidean distance transform for continuous radial position, showing the locations of two spots (pink) within one nucleus. Dark, central: 0. Light, peripheral: 1. Scale bar: 5 µm. C: Diagram of equi-area concentric shell boundaries, showing the locations of two spots (pink) within one nucleus. Note that equi-area shells have square-root spacing relative to radial position itself. Scale bar: 5 µm. D: Maximum projections in XY (top) and YZ (bottom) showing flat, columnar organization of four representative cells. Blue arrows: a “false central” spot that appears relatively far from the periphery in XY but is likely at the very bottom of the cell in XZ. Green arrows: a “true central” spot which appears far from the periphery in XY and is likely in the center of the cell in XZ. Scale bar: 10 µm. E: CDFs showing continuous normalized radial position for all probes tested, separated into panels by chromosome. Position along chromosome is color-coded. F: Correlation between radial positions in a random 1000-cell subsampling at the two alleles of four example loci on chromosome 1, as marked in the figure. There is little to no observed correlation. G: Correlation within a 1000-pair random subsample between radial position at probe 609 on chromosome 4 (x axis) and radial position at multiple adjacent target probes on chromosome 4 as specified (y-axis). Genomic distance between probe centers ranges from 0.25 Mbp (609-610) to 2.28 Mbp (609-618). H: Correlation within a 1000-pair random subsample between radial position at pairs of probes, with genomic distance between probes and chromosome as marked. Kbp: kilobasepairs. Mbp: megabasepairs. Probe pairs used (clockwise from top left): chr1 73.74, chr1 89.91, chr17 142.146, chr1 91.433, chr 17 40.86, and chr 18 186.193.

FISH signals were identified using the Laplacian of the Gaussian method, which performed as well as previously used deep learning segmentation (46) (Fig S1B,C), and centers of gravity were determined for each spot in X and Y as this reduced noise in distance distributions (Fig S1D, see Materials and Methods). The radial position for each probe signal was calculated based on the distance to the edge of the segmented cell nucleus in the DAPI channel (Fig 1B). Alternatively, the nuclear area was binned into concentric shells of equal area, and spots were assigned a radial shell according to their radial position (Fig 1C). Locus positions were determined in 2D projections as previously described (47) rather than in 3D reconstructions since the elongation of signals in the Z direction made it likely that a signal was visible in most Z positions within the nucleus and because the out-of-focus light and slight irregularities in the imaging well made it challenging to approximate the top or bottom of all the cells (Fig 1D). At the length scale used here, 2D analysis of projected images performs as well as 3D analysis for comparisons between two probe pairs or conditions (48). Visual examination was occasionally used to confirm the localization of loci (Fig 1D).

### Radial positioning of genome loci is intrinsically variable

We first examined the population-level radial positioning of each tested probe using an empirical cumulative distribution function (ECDF; Fig 1E). As previously observed, different loci showed distinct localization preferences (Fig 1E; (43)). As entire chromosomes have characteristic radial positions, probes on the same chromosomes showed, as expected, similar distribution profiles, with the overall range of radial positioning preferences being somewhat smaller on chromosomes 17 and 18 than chromosomes 1 and 4 (Fig 1E). The two large regions on chromosome 4 with multiple loci tested per region showed different radial positioning patterns from each other but more consistent radial positioning across the contiguous genomic region (Fig 1E, yellow lines vs blue lines).

We next asked whether the radial position of the two homologs in a nucleus are correlated to determine whether radial position varies intrinsically on the basis of the chromosome or extrinsically on the basis of the cell (Fig 1F). Analysis of 69 loci indicated that the radial position of the two homologs showed little correlation in individual cells (Fig 1F, Fig S2A; maximum adjusted r^2^: 0.040), in line with our previous observation that chromatin interactions were not correlated between homologs (Finn et al. 2019). The position of loci varied independently within a preferred position, consistent with each individual chromosome’s nuclear position being largely independent (Fig 1F, Fig S2A). As expected, adjacent loci on a single chromosome often were strongly correlated at distances up to several million base pairs (Fig 1G, H) even between some loci spaced up to 12 Mbp apart (Fig 1H), but loci spaced by extremely long distances (> 20 Mbp) often lost this correlation (Fig 1H, Fig S3). These observations suggest that large regions within the genome, composed of multiple topological domains, move together relative to the nuclear periphery.

We next asked whether the measured radial position distributions correlated with LADs. For this assessment, we used publicly available DAM-ID sequencing data generated on the same fibroblast cell line (49, 50). We compared the radial position of loci on chromosomes 1, 17 and 18 to the DAM-ID score (Fig S2B), as well as chromatin compartment assignments on the basis of Hi-C data (49, 50). We observed a general trend for the most-peripheral loci to be in LADs, and for peripheral loci to be more likely to be in LADs and B compartment heterochromatin than inter-LADs and A compartment euchromatin (Fig S2B). This trend was especially visible on chromosome 18, and somewhat apparent on chromosome 1, although inter-LAD loci were found at a very wide variety of radial positions (Fig S2B). We did not observe any correlation between LAD status or compartment and radial position on chromosome 17 (Fig S2B). We also examined the enrichment of specific histone modifications (H3K4me3, H3K27ac) in published ChIP-seq datasets (51), and found no correlation between median or most common radial position and chromatin type as defined by either of these two metrics (Fig S2B). These findings are in line with the observation that LADs have been shown to not exclusively be localized to the nuclear periphery but also in the nuclear interior in individual cells (52).

### Interactions between genome loci are enriched at specific radial positions

We next sought to determine whether pairwise interactions between two loci showed a preference for their radial location. To this end, we measured the spatial distance between interacting loci and at the same time determined the radial positions of each signal in the pair (Fig 2). With the exception of genomically proximal pairs on chromosome 4, most of these pairs colocalized rarely and the radial positions of the two loci were frequently quite different. As either locus of an interaction pair could show a preference for localization at a specific nuclear position, we separately compared spatial distances between the two loci to the radial position of each locus in the pair. Visual examination of 2D density plots of radial position versus interaction distance revealed several distinct patterns (Fig 2A). Some locus pairs interacted more commonly at the center of the nucleus (Fig 2A, top panel) whereas others interacted more commonly at the periphery (Fig 2A, bottom panel). These correlations were statistically significant when considered with an ANOVA test (Top panel: slope = 1.38, p = 4.93*10^−8^, Bottom panel: slope = −1.13, p = 1.41*10^−8^).

**Figure 2:**
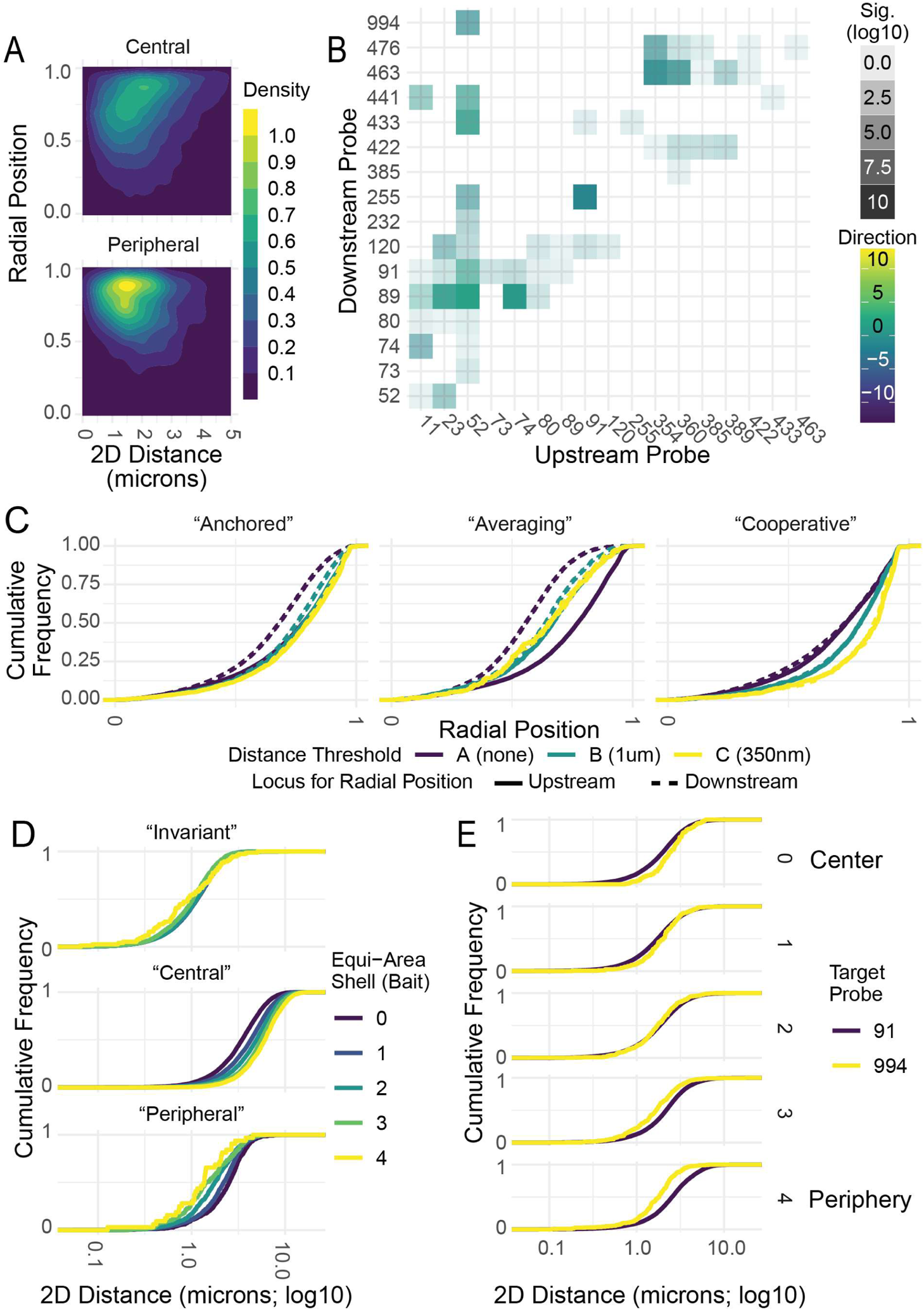
Pairwise associations show preferred radial positions. A: Two examples of 2D density plots showing a positive correlation between radial position and 2D spatial distance (i.e. interactions occur in the center of the nucleus) and a negative correlation between radial position and 2D spatial distance (i.e. interactions occur at the periphery). Data used is a 1000-pair random subsample for each plot. B: Heatmap showing overall correlation between radial position at the upstream locus and distance from the upstream to downstream locus for all pairs on chromosome 1 tested in HFFs. C: Examples of CDFs showing cumulative distributions of radial positions for both spots in pairs with various patterns of enrichment: one “Anchored” pair (chr1 11.52), one “Averaging” pair (chr1 52.89), and one “Cooperative” pair (chr1 52.255). 2D distance threshold between spots is shown in color (purple: farther apart than 1 μm, teal: between 1 μm and 350 nm, yellow: within 350 nm). Solid lines show the radial position of the upstream probe, dashed lines show the radial position of the downstream probe. D: CDFs showing 2D spatial distance on a log scale, color-coded by radial shell of the upstream probe in the pair. “Invariant” (chr17 86.123): no dependence between shell and spatial distance. “Central” (chr17 121.142): more central shells (shell 0) show shorter distances. “Peripheral” (chr1 91.255): more peripheral shells (shell 4) show shorter distances. E: CDFs showing position-dependent partner preference, with 2D distance to locus 52 on a log scale on the x-axis, comparing a peripherally-enriched partner (994, in yellow) to a centrally-enriched partner (91, in purple). Panels show radial shell of locus 52 (0 is most central, 4 is most peripheral).

To comprehensively analyze the relationship of interaction frequency and location, we calculated four statistical parameters for each of the 137 interaction pairs: 1) the direction and 2) significance of correlation between radial position at the upstream probe and distance between the probes, and 3) the direction and 4) significance of correlation between radial position at the downstream probe and distance between the probes. To calculate significance, we used an ANOVA test to determine whether variance in radial position of either locus significantly contributed to variance in distance between the spots. To calculate direction, we used the slope of a least-squares linear regression model to determine whether interactions were more common at peripheral or central locations. Instances where spot-to-spot distance depended on radial position were in fact common (Fig 2B, S2). Of the 137 pairs examined, 58% showed a significant correlation between spatial distance and radial position of at least one spot (p < 0.01) and 22% showed a significant correlation between radial position of both spots (p < 0.01). These correlations occurred at similar rates for enrichment at the edge of the nucleus or enrichment in the center of the nucleus. 45% of pairs interacted more frequently when one of the participating loci was at the edge of the nucleus, and 35% of pairs interacted more frequently when one of the participating loci was in the center of the nucleus. Thus, a dependence on radial position is common, but not universal, and the degree and direction of enrichment appears to be specific to the interaction. We conclude that the frequency of chromatin interactions is often related to their 3D nuclear position.

The 80 locus pairs that showed a correlation with location fell within three distinct groups with regard to their behavior (Fig 2C). First, for 36% of pairs a significant association between radial position and pairwise distance was only present at one locus in the pair. We termed these “Anchored” interactions since an ‘anchored’ spot was found at the same overall radial position irrespective of its distance to its ‘variable’ interaction partner. In these cases, the ‘variable’ spot’s preferred radial position converged on this ‘anchored’ spot’s radial position as distance thresholds were decreased (Fig 2C). Second, 12% of pairs showed “Averaging” interactions, where at large spatial distances, the two loci took on very different radial positions, but the radial position of interacting loci converged on an intermediate pattern (Fig 2C). Finally, 10% of pairs we termed “Cooperative”, as the radial position distributions for two loci were similar at all spatial distance thresholds, but interacting locus pairs occurred at different positions than pairs that were spatially farther apart (Fig 2C). It is worth noting that broad trends were visible between the types of pairs and their association patterns. “Averaging” interactions tended to occur between loci with disparate preferred radial positions, such as those between LADs and interLADs (9/16 pairs) or instances where one locus was most often peripheral and the other most often central (12/16 pairs). Similarly, “Cooperative” interactions tended to occur between two LADs or two interLADs (11/14 pairs) or between loci with similar radial positions (10/14 pairs).

The occurrence of significant, and pair-specific, associations between 2D spatial distance and radial position were confirmed by analysis of equi-area shells (Fig 2D). We observed some interaction pairs with no significant dependence on radial position (top panel, Fig 2D), pairs where shorter distances occurred in more central shells (middle panel, Fig 2D), and pairs where shorter distances occurred in more peripheral shells (bottom panel, Fig 2D). We also observed that a locus can have different behaviors with different interacting partners. For example, probe 52 (chr1:12708720-12865743) tends to be spatially closer to locus 91 (10 Mbp away at chr1:22223363-22394657) when it is central and locus 994 (almost 250 Mbp away at chr1:248125294-248263553) when it is peripheral (Fig 2E). This tendency for one region to have different preferred partners at different radial positions suggests that repositioning a locus from one nuclear subcompartment to another may introduce that locus to a different chromatin environment, rather than pulling its most common interactors along with it (Fig 2E).

### Cell-type specificity and chromatin dependence of preferential interaction sites

Chromatin properties differ between ES cells and differentiated cells. In particular, ESC chromatin is generally more homogeneously decondensed and is hyperdynamic with regards to binding of chromatin proteins (53–55). We took advantage of these differences in chromatin properties in HFFs and H1 human embryonic stem cells (hESCs) to ask whether the observed position dependence of interactions is cell-type specific and affected by chromatin state. To do so, we selected a set of 26 interactions between 9 loci spanning approximately 30 Mbp on the p-arm of chromosome 1, which in HFFs show both “Cooperative” and “Averaging” patterns (Fig 3A, yellow tags; all probes in purple). We comparatively mapped the location and interaction frequencies of these loci in H1 hESCs and HFFs (Fig 3).

**Figure 3:**
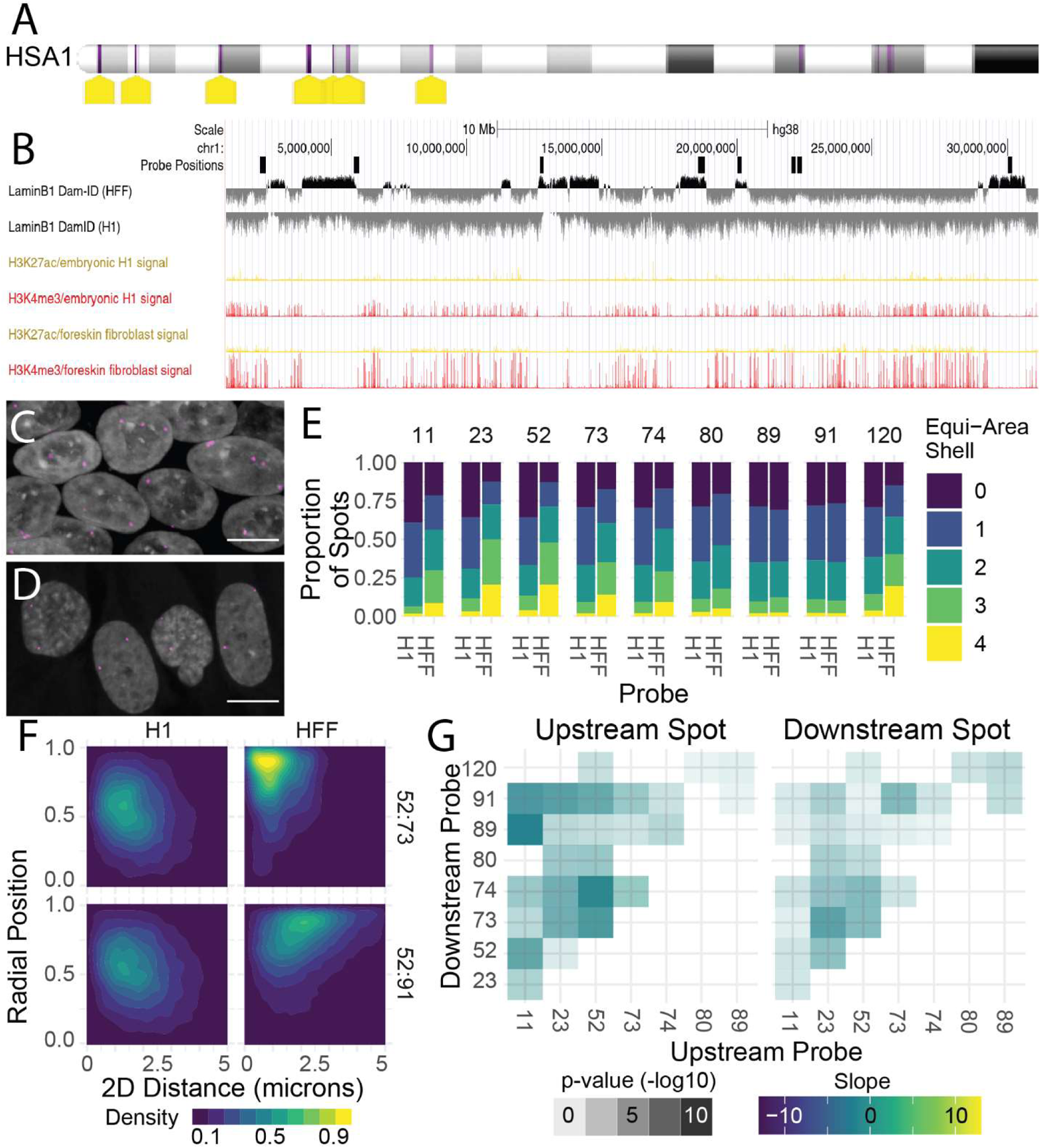
Embryonic stem cells show centralization of probes but peripheral trend for interactions. A: Schematic location of all probes tested in H1 embryonic cells on chromosome 1. Yellow tags: probes stained in H1 cells; purple bars: all probes stained in HFF cells. B: UCSC Genome Browser track showing LAD position and chromatin type for a region on the P arm of chromosome 1 in H1 hESCs (embryonic H1) and HFFs (foreskin fibroblast). C: Representative field of H1 human embryonic stem cells: DAPI staining (gray) and FISH (pink). Scale bar: 10 μm. D: Representative field of HFFs: DAPI staining (gray) and FISH (pink). Scale bar: 10 μm. E: Bar charts showing general centralization of all probes in this region in H1 cells relative to their position in HFF cells. F: 2D density plots showing radial position of one locus (locus 52) as a function of 2D spatial distance to a downstream locus (locus 73 or 91 as noted) in both HFFs and H1 hESCs. Data used is a 1000-pair random subsample for each plot. G: Heatmaps showing correlation between radial position of upstream probe and pairwise spatial distance between two probes in H1 cells.

Publicly available sequencing data confirmed the expected chromatin differences between these cell types at this region (Fig 3B). In HFFs, the chromosome 1 region contained seven large LADs and eight additional regions of positive Dam-ID signal that were too small to be classified as LADs (Fig 3B). In contrast, in H1 cells the same region was uniformly inter-LAD (Fig 3B). Similarly, the histone modification landscape of this region is markedly different in H1s than in HFFs (Fig 3B). HFFs show regions of enrichment of H3K27 acetylation and especially H3K4 methylation in inter-LADs and depletion of these marks in LADs, while in H1s, both marks are more uniformly depleted (Fig 3B).

We imaged and analyzed at least 500 H1 hESCs for each locus pair for a total of 153,204 analyzed pairs with a median of 4,572 signals per pair (Table S2). In line with the more homogenous chromatin features in H1 cells, all loci tested were centrally located in H1 cells versus their frequently more peripheral localization in HFFs (Fig 3C-D). Complementary equi-area shell analysis of the entire imaging dataset revealed that while in HFFs many of the loci in this region are commonly found in shells 3 and 4 (Fig 3E), all loci tested in this region have a preference for shells 0 and 1 in H1 cells (Fig 3E).

Similar to HFFs, in 2D density plots pairs of loci showed patterns of enrichments at certain radial positions in H1 cells (Fig 3F). Strikingly, in H1 cells, all interaction events showed the same trend. Interactions were slightly more common at peripheral positions, regardless of whether they were enriched at peripheral or central positions in HFFs (Fig 3F). We quantitated the relationship between location and spatial distance at every tested pair in H1 cells as we did in HFFs and confirmed this more consistent trend in H1 cells. While 58% (80/137) of pairs showed significant correlation between pairwise distance and radial position in HFF cells, 85% (22/26) of pairs showed significant correlation between distance and radial position in H1 cells (Fig 3G). This difference is significant by a hypergeometric test (p = 1.84 * 10^−3^). Thus, in every instance, and although the probe pairs chosen were selected due to the variety of patterns they exhibited in HFF cells, associations between two probes in H1 cells were almost always more likely to occur at the edge of the nucleus than the center. These differences in chromatin features in HFFs and H1 hESCs, and the correlated changes in the nuclear organization of loci and interacting events, suggest that in genome regions with uniform euchromatic structure there is a preference for interactions to occur at the nuclear periphery.

### Cell-type-specific enrichment of interactions by decreased mean distance or increased variability

The observed increased homogeneity in radial position at these loci in stem cells compared to fibroblasts is consistent with higher homogeneity in genome folding at these loci in stem cells. Examination of the chromosome 1 region in Hi-C maps from publicly-available data indeed confirms areas of enrichment and depletion of interactions in HFFs compared to H1s (Fig 4A). We calculated distance distributions from imaging data to determine the nature of cell-type specific differences in interaction frequency and 2D distance. Overall, the range of average spatial distances was similar between HFFs and H1s. Median distances in both H1s and HFFs ranged from 1.1 µm to 1.9 µm, and as a population were not significantly different (Fig 4B; t-test p = 0.057). After correction for the decreased nuclear size of H1 cells, 2D spatial distances captured a larger proportion of the nuclear area in H1 hESCs than in HFFs (Fig 4C; t-test p = 5.98*10^−4^). A pronounced difference between the two cell types was, however, that in H1 cells, the 2D spatial distance distributions showed an almost perfect dependence on genomic distance, unlike in HFFs (Fig 4B; r^2^ 0.94 in H1s, 0.41 in HFFs). Thus, while the total range of spatial distances may be independent of cell-type, and chromosomes in hESCs may occupy larger proportions of nuclear space, chromatin organization of this region in H1 cells appears to depend almost entirely on genomic distance, whereas in HFFs specific pairwise interactions are often enriched or depleted in relative to expectations based on the genomic distance separating them.

**Figure 4:**
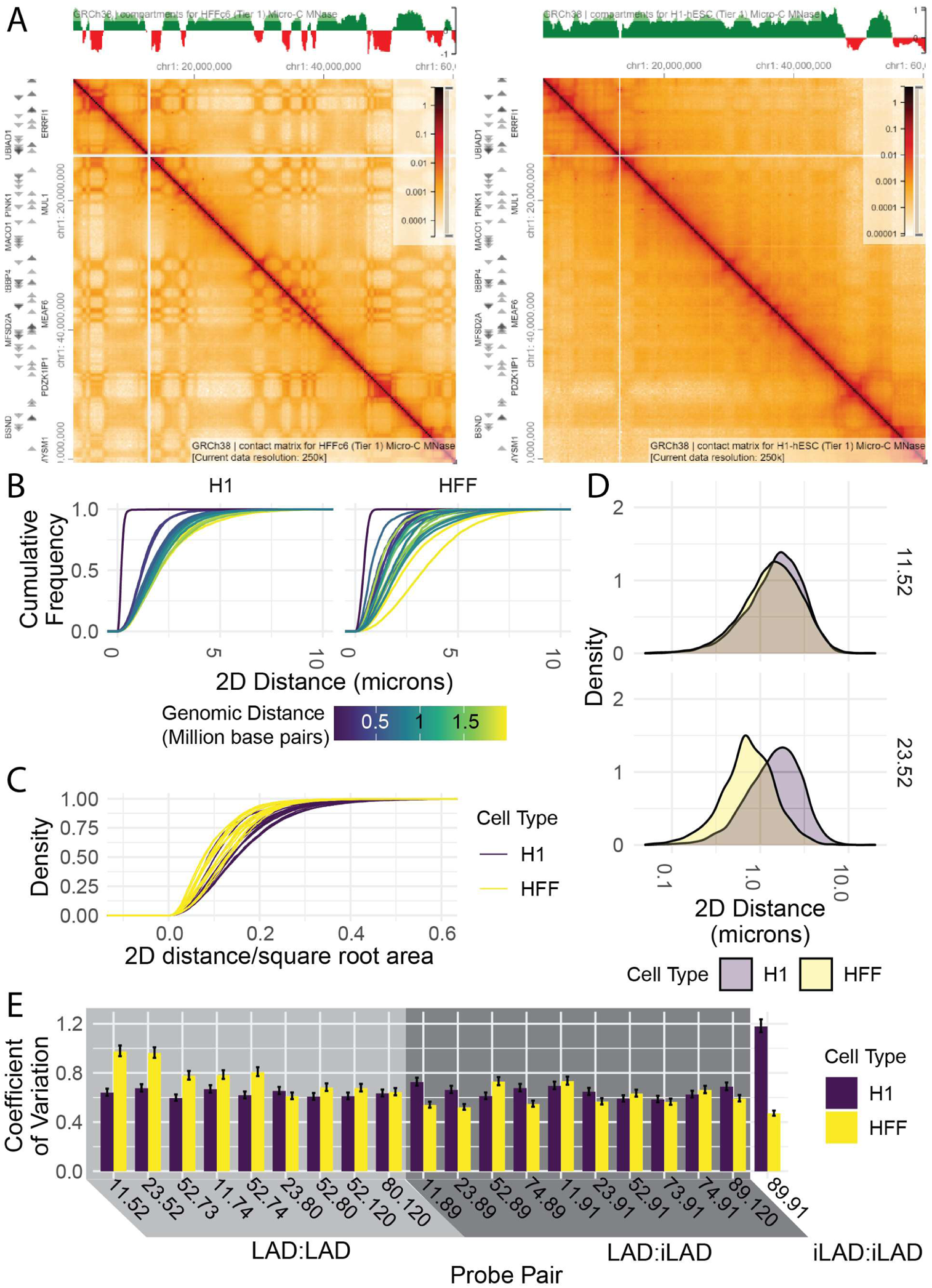
Cell-type-specific pairwise associations have higher coefficients of variation. A: Micro-C plots and compartment scores for studied region on the p-arm of chromosome 1 in HFF-c6 and H1 hESC cells. B: CDFs of 2D spatial distances between probes, color-coded by genomic distance, in H1 and HFF cells. C: CDFs of 2D spatial distances normalized to nuclear size (square root of area), color-coded by cell type. D: Representative PDFs showing 2D spatial distances between probes, color-coded by cell type, for two selected probe pairs (top panel: chr1 11.52; bottom panel: chr1 23.52). E: Bar graphs showing coefficient of variation in 2D spatial distance of a 1000-pair random subsample, color-coded by cell type.

Cell-type-specific enriched associations could either arise by a decrease in mean distance or, alternatively, via an increase in variability in spatial distance. To discriminate between these two possibilities, we compared distance distributions of HFF-depleted and HFF-enriched interaction pairs. Statistically significant changes in mean distance and variability were very common in these loci. 6 of 9 HFF-enriched pairs showed a statistically significant decrease in mean distance (t-test, p < 0.05 after Bonferroni correction), and 6 of 10 HFF-depleted pairs showed a statistically significant increase in mean distance (t-test test, p < 0.05 after Bonferroni correction). Similarly, 4 of 9 HFF-enriched and 5 of 10 HFF-depleted pairs showed a statistically significant change in variability (asymptotic test for equality of coefficients of variation, p < 0.05 after Bonferroni correction).

When comparing changes in mean distance and variability of individual pairs, we find that one HFF-enriched interaction and two HFF-depleted interactions showed altered variability but not altered distance. Colocalization between probe 11 (chr1: 2370451-2570719) and probe 52 (chr1: 12708718-1286743), separated by approximately 10 Mbp, is more likely to occur in HFFs than in stem cells (5.2% vs 4.0% within 350 nm; 0.5% vs. 0.2% within 100 nm; p < 0.005 by a two-sample proportion test). However, the mean spatial distance between the two cell types is very similar (Fig 4D; 1.80 µm in fibroblasts vs 1.85 µm in stem cells, p = 0.0183 by a t-test). This is made up for by an increase in variability (Fig 4D; standard deviation 1.58 in fibroblasts and 1.22 in stem cells).

Similarly, in five of nine HFF-enriched pairs and five of ten HFF-depleted pairs, changes in mean distance alone, without a significant change in variability, were sufficient to drive enrichments. This is the case for colocalization between probe 23 (chr1: 5829955-6030168) and probe 52 (chr1: 12708718-1286743), separated by about 7 Mbp. Colocalizations within 350 nm or 100 nm were substantially more common in HFFs than in stem cells (13% vs. 3.8%, p < 2.2*10-16 for 350 nm; 0.89% vs 0.27% for 100 nm, p < 0.0005 by a two-sample proportion test). The mean distance between probes was substantially shorter in HFFs than in H1s (Fig 4D; 0.73 µm in fibroblasts vs 1.52 µm in stem cells; p = 9.42*10^−71^ by a t-test), but HFFs exhibited a lower standard deviation (0.81 for HFFs vs 1.17 for stem cells).

While these results point to distinct behavior of individual locus pairs, it is worth noting that in both examples, the end result is an increased coefficient of variation that was statistically significant (p < 1*10^−4^ in both cases by the asymptotic test). In fact, this trend for an increased coefficient of variation of spatial distance at HFF-enriched locus pairs in HFFs was the most consistent feature of enriched associations (Fig 4E). In every case where an interaction was enriched in HFFs, the coefficient of variation in HFFs was also higher; in half of them it was statistically significant (p < 0.05 after multiple hypothesis testing, see Materials and Methods) (Fig 4E). In the case of interactions that were relatively depleted in HFFs, the differences in coefficient of variation were more mixed; while many were statistically significant after correction, some pairs showed higher variability in H1 cells and others showed higher variability in HFF cells (Fig 4E). This behavior is consistent with a model for long-range heterochromatin organization whereby both restricting the spatial distance or increasing the mobility of loci can alter interaction frequency.

Our analysis in H1 cells not only shows that genome structure in H1s is overall more uniform than in HFFs, but also that the non-uniformity in HFFs may drive specific chromatin-chromatin associations.

## Discussion

Genome mapping methods have found that both the position of a locus relative to the edge of the nucleus and the position of a locus relative to other loci are non-random (1). Despite the emergence of non-random patterns, genome organization is also characterized by extensive single-cell variability. Chromosome positioning, and radial positioning of specific genes, has long been characterized as probabilistic rather than deterministic (56) and chromatin interactions between genomic sites are marked by considerable intrinsic variability (35, 37). However, studies comparing the effect of the nuclear position of genomic loci on their propensity to undergo chromatin interactions in endogenous systems have been lacking. Here we used high-throughput imaging to simultaneously determine the radial coordinates and the frequency of interaction for 137 chromatin interaction pairs between 69 loci on four chromosomes in HFF cells and 26 interactions between 9 loci on chromosome 1 in H1 human embryonic stem cells to ask whether the position within the nucleus of loci affects their interaction frequency. We find that the position within the nucleus is associated with differential pairing frequencies, and that loss of preferred localization within the nucleus is related to decreased rates of enriched associations as well as decreased variabilities.

Most of the locus pairs we tested showed some dependence between nuclear position and interaction frequency, suggesting an interrelation between genome folding and nuclear organization. Overall, we find that the periphery of the nucleus is a more favorable environment for pairwise associations than the nuclear interior. This is particularly prominent in embryonic stem cells, as while overall the chromatin landscape of the region we tested was euchromatic and centrally located, every tested locus pair was more likely to interact when at the edge of the nucleus. However, it is also apparent among loci not excluded from the nuclear periphery in HFFs, as loci characterized as inter-LADs which nonetheless could be found at the nuclear periphery interacted more frequently when peripheral. These observations suggest that the nuclear periphery facilitates especially long-range interactions.

A simple explanation for this observation is that the periphery is characterized by increased chromatin density, which would shorten all local distances. Two observations argue against this interpretation. First, the density and segregation of peripheral heterochromatin increases during differentiation (57), and the periphery in stem cells is marked by lower levels of A-type lamins and heterochromatic marks (58, 59), so the expectation would be that a chromatin-density-mediated effect would be stronger in HFF cells than H1 hESCs, which is contrary to what we observe. Second, a chromatin density effect would be more prevalent at short-range associations, which reliably move together into the denser environment of the nuclear periphery. In contrast to this prediction, we find trends for short-range as well as long-range associations, and the strongest trends in fibroblasts are for long-range associations between loci whose radial positions correlate only poorly. Instead, it seems more likely that the observed preference for peripheral localization of interactions in embryonic stem cells is largely due to geometric effects, whereby short-range movements of two loci brought together early in G1, when higher order genome organization is established, cannot move loci far apart at the edge as easily as they can in the center. In fact, slower movement of loci tethered to the nuclear periphery has been observed (60). The relative immobility of the interaction partners may increase their likelihood of spatial proximity throughout the rest of the cell cycle. In this model, the same geometric constraints – and the tethering of regions to the nuclear lamina – act to reduce the ‘search volume’ for particular long-range interactions, thus increasing their rates of interaction as we observe in fibroblasts.

We find that the correlations between nuclear organization and genome folding are generally stronger in fibroblasts compared to hESCs, and that the directionality of these correlations is more variable in fibroblasts than in hESCs. These trends coincide with differences in chromatin structure between fibroblasts and hESCs, pointing to a role of nuclear subcompartmentalization in chromatin folding. Stem cells have a unique, generally more open and globally permissive chromatin environment with overall increased levels of euchromatic marks (54) and reactivation of otherwise stably silenced transposable elements (61). Significant and global rearrangements of the heterochromatin compartment occur during differentiation (57). In line with these properties, we find that in stem cells there is a consistent effect of location on most interaction pairs, whereas in fibroblasts we observe pair-specific effects, suggesting that the trend for specific pairs to interact at specific locations within the nucleus may depend on their chromatin type. In stem cells, which do not show differential nuclear organization at the loci analyzed, preferential pairing between loci is also lost. Furthermore, even within fibroblasts, we observed that enriched associations between LADs tended to occur, perhaps unsurprisingly, at the edge of the nucleus, and that in the relatively rare instances where normally peripheral loci were found in the center of the nucleus, they were more likely to be in spatial proximity to genome loci with which they rarely interacted in the overall population. Thus, it seems reasonable to conclude that a strongly heterotypic chromatin landscape arising during differentiation allows the segregation of frequent interacting pairs, but flexibility in nuclear organization as observed in stem cells allows loci to sample the entire range of nuclear positions and interacting partners, thus maintaining plasticity.

Based on the higher level of compartmentalization and higher degree of heterochromatin in fibroblasts, one might expect that fibroblast-enriched pairing events would show decreased variability in their separation distance. In contrast we find increased variability of highly interacting loci in fibroblasts. We observe increases in colocalizations concomitant with decreased mean distance without a change in variation and with increased variation without a change in mean distance. While there are many systems in which enrichment in chromatin interaction frequency does not correlate with mean distance (62), our observations are among the first to directly show that increased variability in distance can lead to enrichment of interactions. This behavior could be due to increased local mobility of the interacting loci, enabling them to scan a larger 3D space for potential interaction partners, for instance when enhancer elements become more mobile upon activation (63). Alternatively, it could be because in a differentiated nucleus with bigger differences between nuclear subcompartments, there are more distinct options for chromosome conformations. It is tempting to hypothesize that chromatin segregation, in addition to separating heterochromatin and euchromatin, may also alter variability and plasticity in chromatin interactions.

There are three technical caveats to our study. First, our use of 2D rather than 3D measurements. second, the relatively small size of the probe sets relative to the genome, and third, the fact that our work is correlative rather than mechanistic. We do not believe that either of these limitations invalidates our findings. 2D measurements on projected images were used for practical reasons so as to reduce computation time and increase throughput. Prior analysis demonstrated very limited loss of accuracy of distance measurements on the length scales analyzed here (48). In addition, errors introduced by 2D approximations would cause us to underestimate the rates of true peripheral localizations and thus make it more difficult to detect correlations between spatial distance and nuclear position as we have found here. The correlation we observed between LAD status or chromatin marks and radial positioning was indeed not strong and might be strengthened by segmenting nuclei in 3D. Similarly, for pairs where we did not see a dependence between radial position and pairwise distance, we cannot rule out the possibility that with 3D positions such a dependence would be detected. While it is therefore possible that we missed some dependencies that truly exist in 3D data, it is not likely that the dependencies we observed are not present. Furthermore, although our probe set is limited, the loci cover a wide range of chromatin features including location in diverse chromatin environment, short- and long-range interactions and locations in gene dense and gene poor genome regions. More comprehensive analysis should be possible using whole-genome-imaging techniques (64). Finally, the differences we observe between H1 hESCs and HFFs are correlations, and proper mechanistic studies of isogenic cells – such as altering the structure of the nuclear periphery using genetic techniques in differentiated cells, or specifically targeting individual model loci to different nuclear compartments and examining the effects on pairwise distances – are required to flesh out the true mechanisms driving the differential organization observed here.

Taken together, our results uncover an interdependence of chromatin interactions and radial positions in the cell nucleus. Our findings point to a close interplay of local chromatin folding and higher order nuclear organization and they highlight the need of using a multi-parametric and, ultimately, a multi-omics approach to studying nuclear architecture.

## Materials and Methods

### HFF culture

HFFs were grown and plated for imaging exactly as described previously (37).

### H1 ESC cell culture

H1 human embryonic stem cells were grown on Matrigel (Corning #354277) in complete mTeSR media (StemCell Technologies). They were split every five days or when colonies began to merge, at a ratio of 1:10-1:12 depending on density. Matrigel was aliquoted into portions equivalent to ¼ the lot-specified dilution factor, sufficient to dilute into 6.25 mL for plating all wells in a 6-well plate. Matrigel plates were made fresh at each split: one aliquot of Matrigel was diluted with 6.25 mL DMEM-F12 on ice, and 1 mL diluted Matrigel was added to each well of an ice-cold 6-well dish. The dish was rotated and shaken to make sure the entire surface of the dish was covered with Matrigel, and allowed to solidify at room temperature for at least 30 min. The supernatant was then removed and replaced with 2 mL room temperature mTeSR media. To split, cells were first washed with PBS and then treated with 1 mL ReLeSR cell dissociation reagent. The majority of the dissociation reagent was gently aspirated from the cells and they were placed in a humid incubator for 6 min. After this incubation, fresh mTeSR media was placed in the well and colonies were lifted by vigorously tapping the dish for 45s. The resulting cell suspension was pipetted once through a 1 mL serological pipette to break down colonies and transfer them into a 15 mL conical tube for passaging. The well was then washed with an additional 1 mL mTeSR media, and an appropriate amount of cell suspension was added to each well of the prepared plate (200 µL for a 1:10 split). The plate was shaken to spread cells evenly before being returned to the incubator. Between splits, media was replaced with 2 mL fresh room temperature complete mTeSR each day.

For FISH experiments, 384-well plates were coated with Matrigel exactly as described above, with a volume of 25 µL diluted Matrigel per well of a 384-well plate. To plate as colonies, 800 µL suspended colonies were diluted 1:10 with 7.2 mL mTeSR, and 50 µL of this diluted cell suspension was added to each well of the prepared 384 well dish. Cells were grown as colonies for three days before fixation.

To plate as a single cell suspension, cells were rinsed once with PBS and incubated in 1 mL ReLeSR/well of a 6-well at 37°C for 8 min. Cells were lifted and dissociated by pipetting with a 1 mL micropipette three times and added to a tube containing 1 mL DMEM/F-12 to neutralize the ReLeSR. Wells were washed with 1 mL fresh DMEM/F-12 for a total of 3 mL cell suspension per well. Cells were then pelleted by spinning for 5 min at 300 x g at room temperature, and media was replaced with 5 mL complete mTeSR media supplemented with 1X ReVitaCell ROCK Inhibitor (Thermo Fisher). Cells were gently, but completely, resuspended by tapping the tube as media was being added and subsequently pipetted twice with a 1 mL micropipette. 100 µL cell suspension was plated per well of a 384-well plate, and cells were spun for 1 min at 300 x g at room temperature to settle and spread. Monolayer cells were grown overnight and fixed the next day. Experimental results were comparable in both cases and as such data were pooled between dispersed and colony-grown measurements.

### DNA FISH

DNA FISH was done as described (65).

### Imaging

Imaging of HFFs was done as described (65).

H1 colonies were imaged in an automated fashion on a Yokogawa CV7000 microscope. To automatically find H1 colonies, we used Wako’s SearchFirst platform (Wako Software Suite). Each well in the plate was initially imaged in the blue channel (Ex: 405 nm, Em: 445/45 nm) at 4x magnification. Colonies were segmented in these images using KNIME, and a uniform tiled grid of all positions in the well within colonies was determined. Of these, 20 positions were chosen at random to image in four colors at 60X. Imaging proceeded as for HFFs on the Yokogawa CV7000 microscope, with the exception of the height of the z-stack, which was increased to 15 µm in order to capture the entire nucleus.

### Computational Analysis

Image processing, including nuclear segmentation, spot segmentation, calculation of the center of gravity in three dimensions and radial position were performed in Python using this Jupyter Notebook (https://github.com/elfinn/cell-and-spot-segmentation). In brief, images were maximally projected prior to segmentation of nuclei using CellPose, a deep-learning based model for segmentation of nuclei including those which touch or have slight overlaps in the 2D maximum projection (66). Centers of gravity in Z were calculated at this step for each pixel in the image where possible. Individual nuclei were cropped in each channel and radial positions for each pixel within each nucleus were pre-calculated using an exact Euclidean distance transform and normalized such that fully central pixels had radial position 0 and edge-pixels had radial position. Nuclei were also binned into five concentric shells of equal area, and spots were assigned to whichever shell their central pixel fell. Within each nucleus, spots were identified using the Laplacian of the Gaussian, which while not quite as sensitive at segmenting spots as a previously trained neural network (Fig S1B, (46)) nonetheless resulted in accurate, and in fact indistinguishable, distance distributions (Fig S1C).

We determined Cartesian coordinates for all spots as centers of gravity as previously applied to localizations of small probes (67), which resulted in a reduction of noise in the distance distributions and a shift in median distance for positive colocalization controls (same probe stained with two colors) from 323 nm to 111 nm (Fig S1D). For each spot, we recorded radial position at the pixel-level spot center and center of gravity calculated in X, Y, and Z where possible for the region defined by half maximum signal normalized to cellular background around each spot center. Subsequent pairwise distances were calculated and statistical analyses performed in R, as described below.

To select cells in which FISH staining and segmentation had proceeded robustly, we considered only those cells which met all the following criteria: 1) At least two spots in each relevant channel, 2) the same number of spots in each relevant channel, 3) coming from a well in which at least 60% of nuclei have exactly two spots segmented. All pairwise distances between spots in different colors were calculated, and minimum distances on a per-green-spot basis (or per-red-spot in the case of comparisons between red and far-red probes) were selected. This final dataset included a list of all spots considered, with X, Y, and radial coordinates (a total of approximately 1,643,000 spots), as well as a list of all minimum pairwise distances along with X, Y, and radial coordinates for both spots in the pair (a total of approximately 1,116,000 pairs).

To map probed loci to previously published sequencing data: We used Lamin-B1 Dam-ID-seq data and Compartment scores generated from Micro-C data in the HFF-c6 and H1 cell lines from the 4D Nucleome consortium (49, 50) (accession numbers: 4DNESZJMHC3O and 4DNESKOBVYIY respectively for micro-C and 4DNESXZ4FW4T and 4DNESXKBPZKQ respectively for LaminB1-DamID-seq) as well as ChIP-seq data from ENCODE (51, 68) (ENCSR814XPE and ENCSR000ANP for H3K4me3 and H3K27ac respectively in H1 hESCs, and ENCSR639PCR and ENCSR510VXV for H3K4me3 and H3K27ac respectively in HFF-c6 cells). In each case, datasets were cross-referenced with our probe sets using the UCSC table browser (https://genome.ucsc.edu/cgi-bin/hgTables). In cases where a BAC probe overlapped multiple annotated features in the dataset, the median score was used. Genomic distances between pairs of loci were calculated from center to center, and distance to the nearest annotated LAD was calculated from edge to edge.

To ensure homogeneity in the sample size across all locus pairs tested, which was essential for some analyses such as ANOVA tests or 2D density graphs, we performed random subsampling with replacement to generate a uniform dataset with 1000 spot pairs per locus pair. When a graph or test was done on subsampled, rather than raw, data, it is indicated in the text. Graphs were generated using ggplot2 (69).

R markdown files to calculate spot distances, perform subsequent analyses, and generate figures are available at https://github.com/elfinn/radpos-r-code.

## Acknowledgements and Funding Sources

This work was supported by funding from the Intramural Research Program of the NIH, the NCI Center for Cancer Research.

## Figure Legends

**Figure S1:**
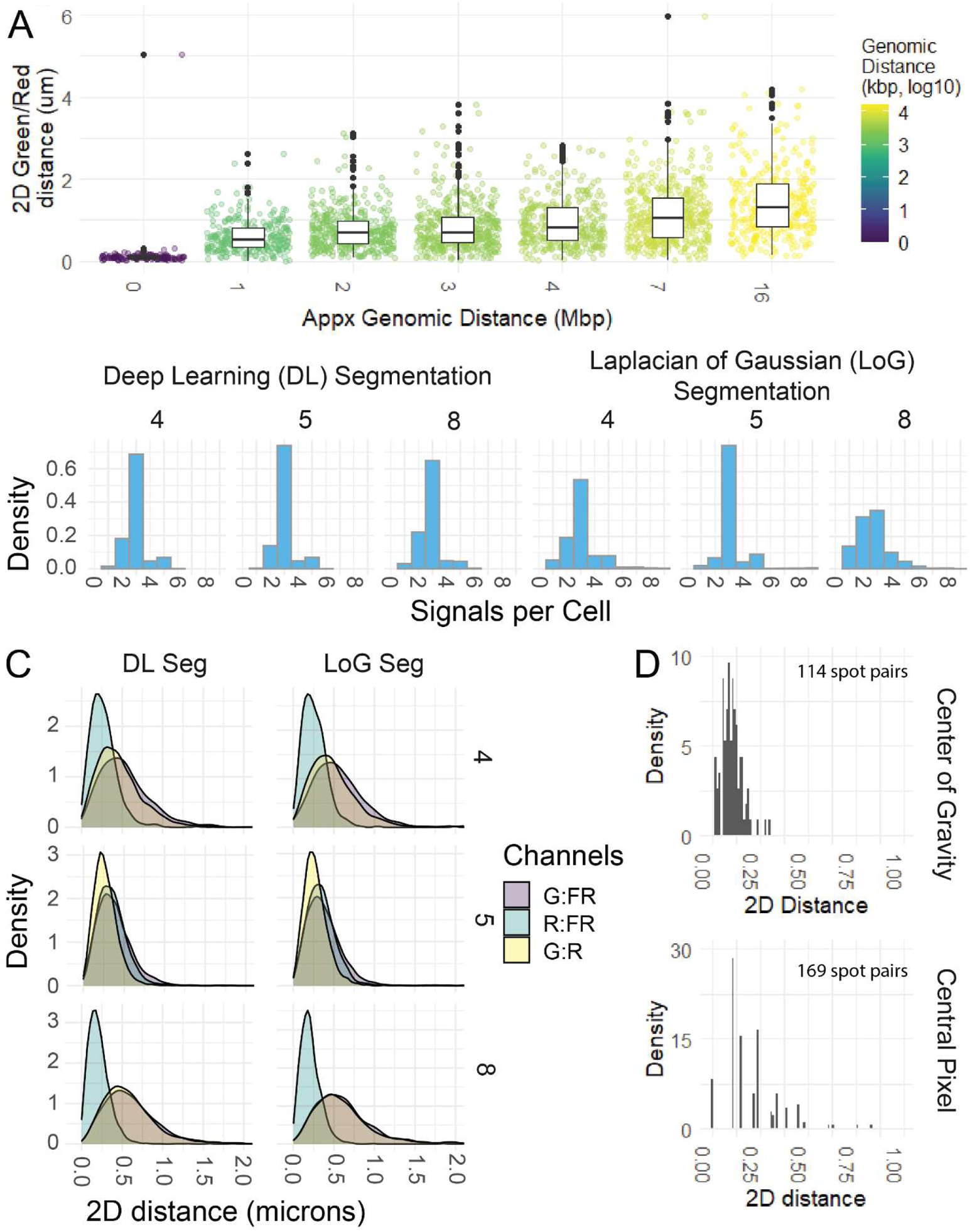
Validation of imaging pipeline. A: Box and jitter plots showing distribution of distances for a selection of probes pairs including a costained control locus (Distance 0) as well as several probe pairs at genomic distances up to ~16 Mbp. Center-to-center genomic distance is color-coded. B: Comparative spots per cell for spots segmented from three representative wells with a deep-learning based published model (46) as compared to and traditional Laplacian of Gaussian-based segmentation. Wells selected for a breadth of FISH quality, from very high signal to noise (well 5) to borderline (well 8). C: Spot-to-spot distance distributions calculated from the spots segmented in (B). Color-coded by pair of channels within the well. D: Spot-to-spot distances for the 120:120 costained locus with spot positions assigned by center of gravity (top) or central pixel (bottom).

**Figure S2:**
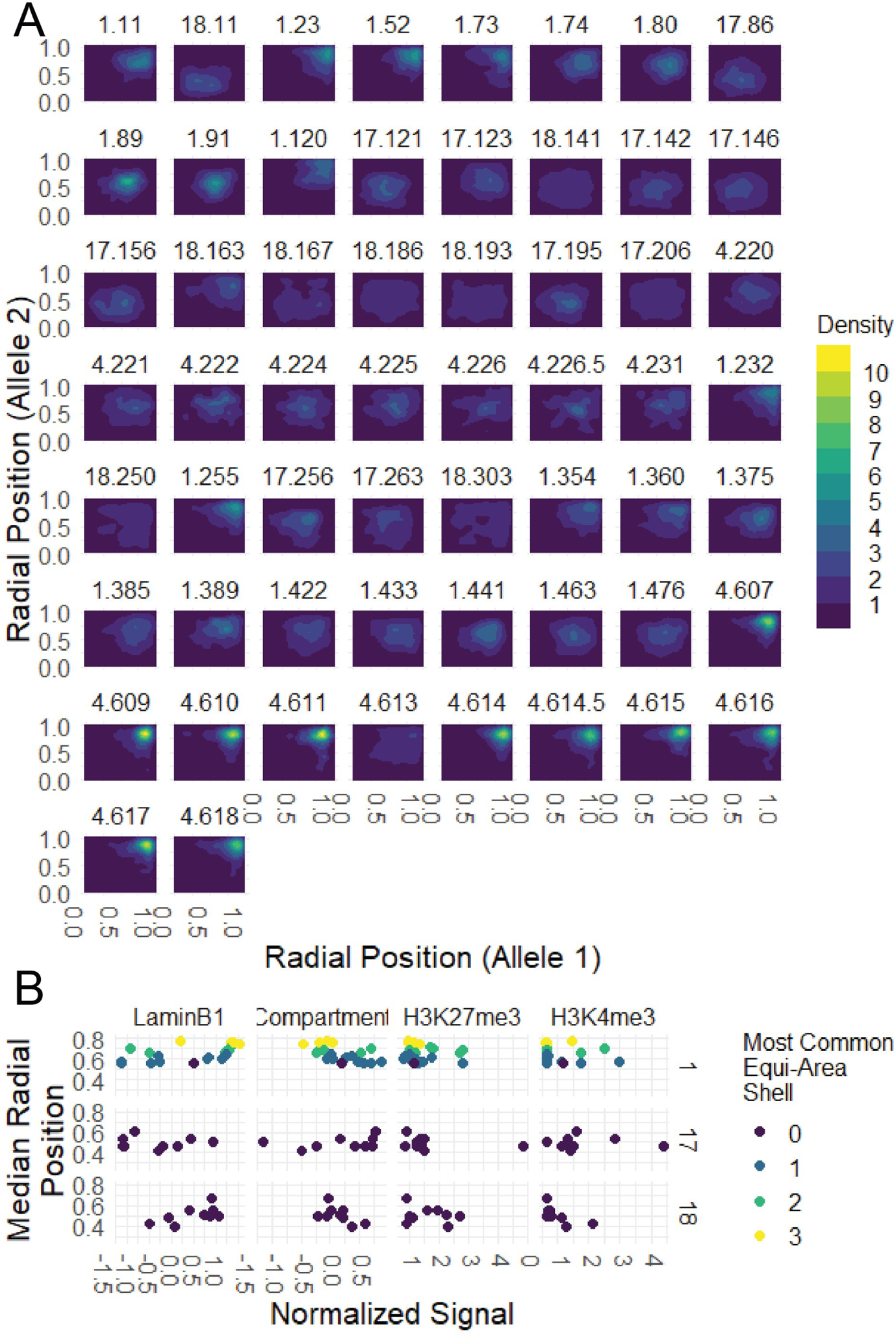
Intrinsic variability in radial position at all tested loci and correlations with sequencing data. A: 2D Density plots showing radial position at one homolog (arbitrarily selected) on the x-axis and radial position at the other homolog on the y-axis. Probe and chromosome as marked. To keep color scheme consistent between panels, spots were randomly subsampled. B: Scatterplots showing median continuous normalized radial position vs. sequencing metrics: LaminB1 enrichment by Dam-ID, Chromatin Compartment in Micro-C, H3K27me3 and H3K4me3 signal in ChIP-seq. Color-coded by most common radial shell.

**Figure S3:**
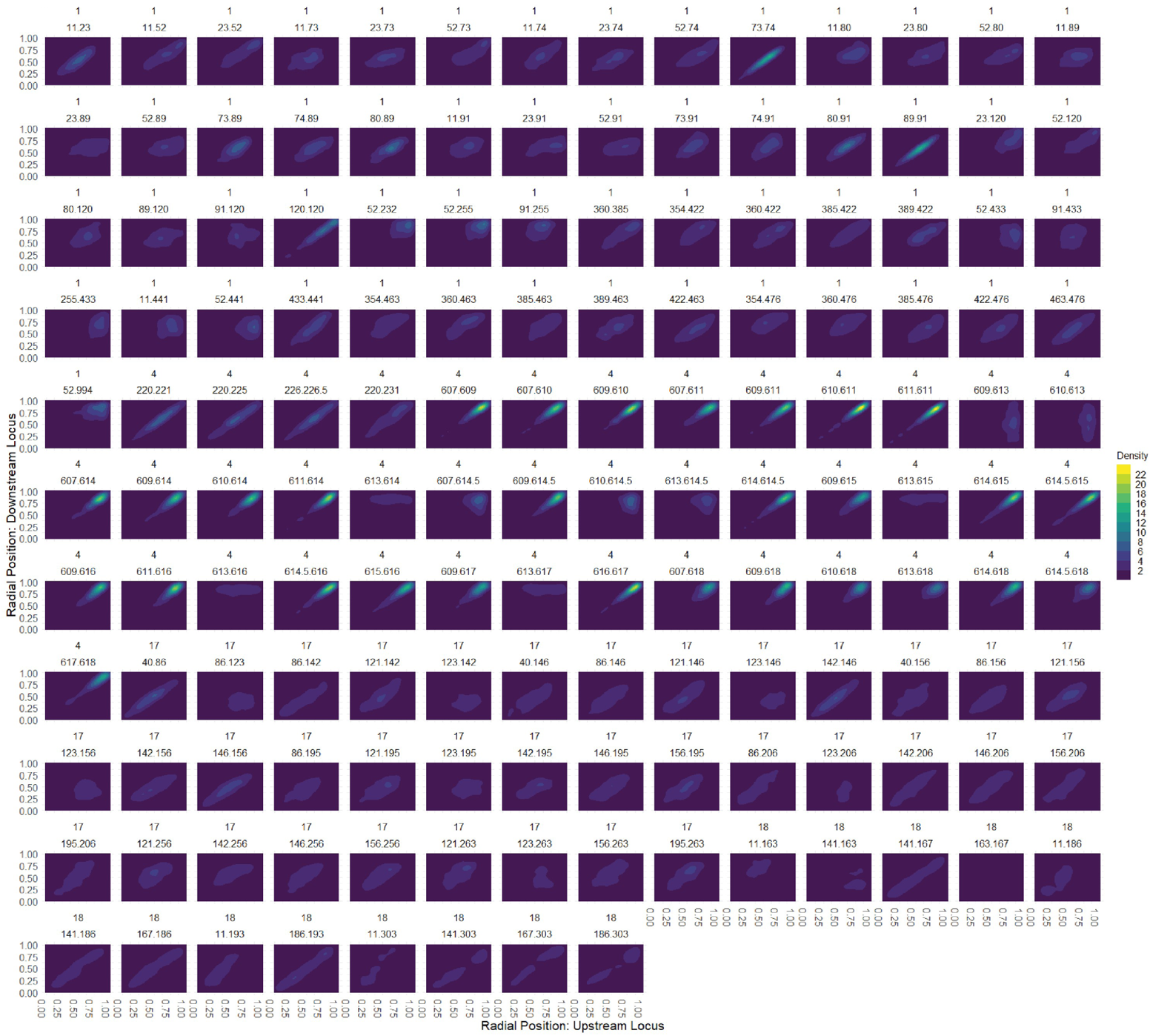
Correlation between radial positions at all probes tested. 2D Density plots showing radial position at one locus in a pair on the x-axis and the other locus on the y-axis. Probe pair and chromosome as marked. To keep color scheme consistent between panels, spot pairs were randomly subsampled.

**Figure S4:**
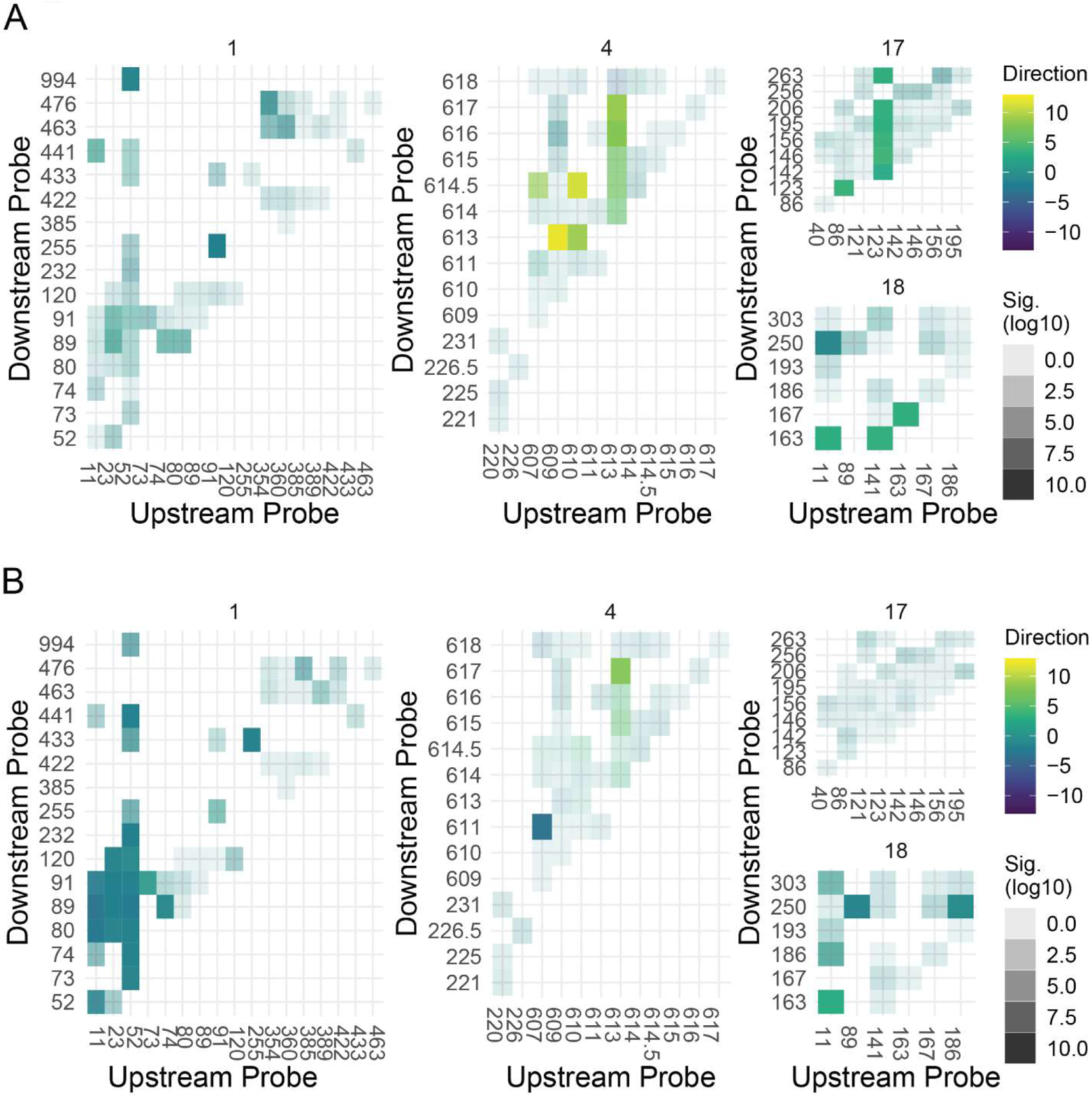
Heatmaps for direction and significance of dependence on radial position for all pairs in HFF cells. Heatmaps for each pairwise interaction on a chromosome, with probe number (approximate genomic position) on both x and y axis. Intensity of color (alpha) is significance (as −log10(p-value) in the ANOVA test). Color is direction (as slope of line of best fit).

**Figure S5:**
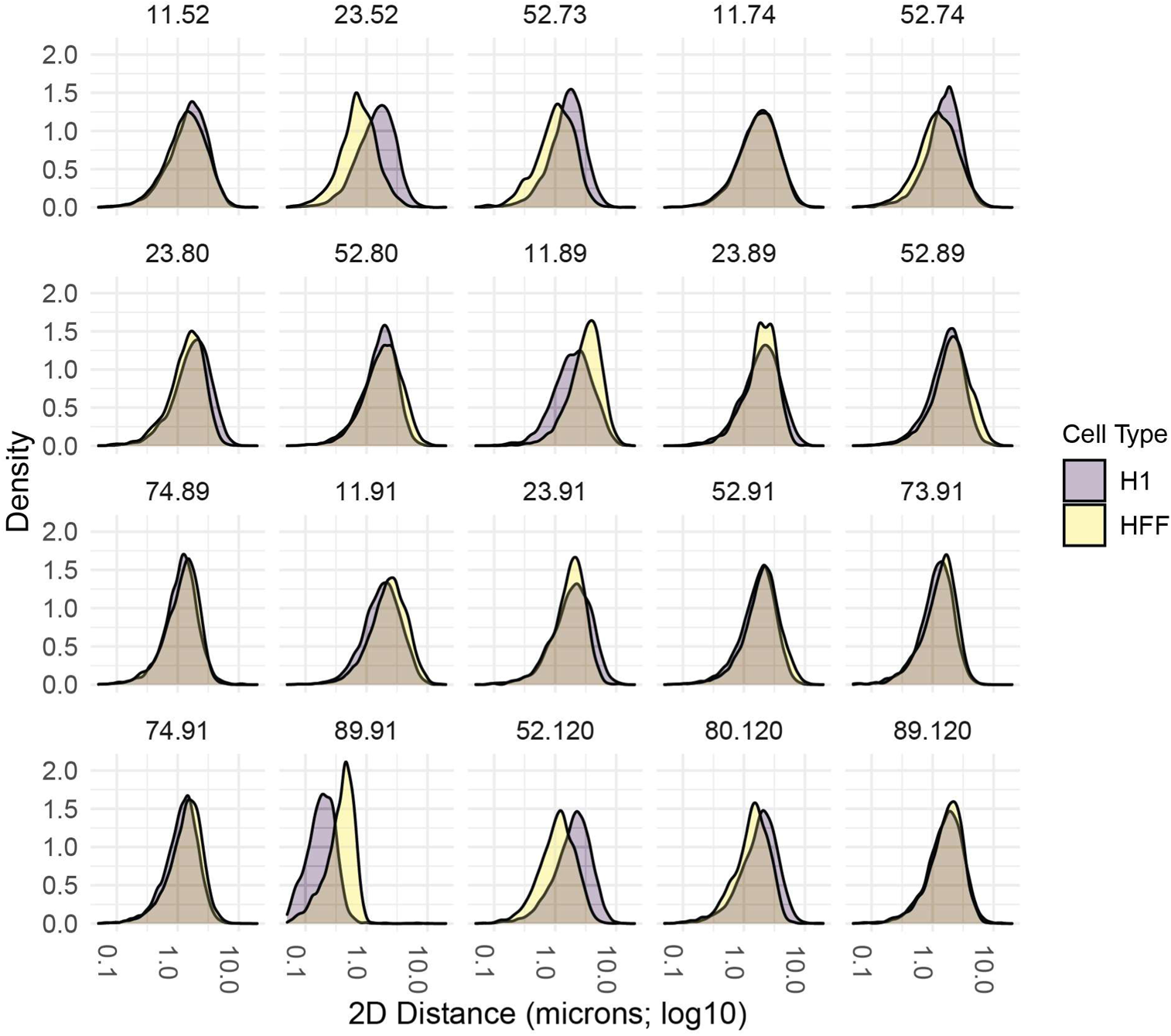
Differences in chromatin folding between HFF and H1. A: Probability density functions showing 2D spatial distance distribution for all pairs examined in both HFFs and H1s. Pair is as marked, color-coding by cell type.

**Table S1:**
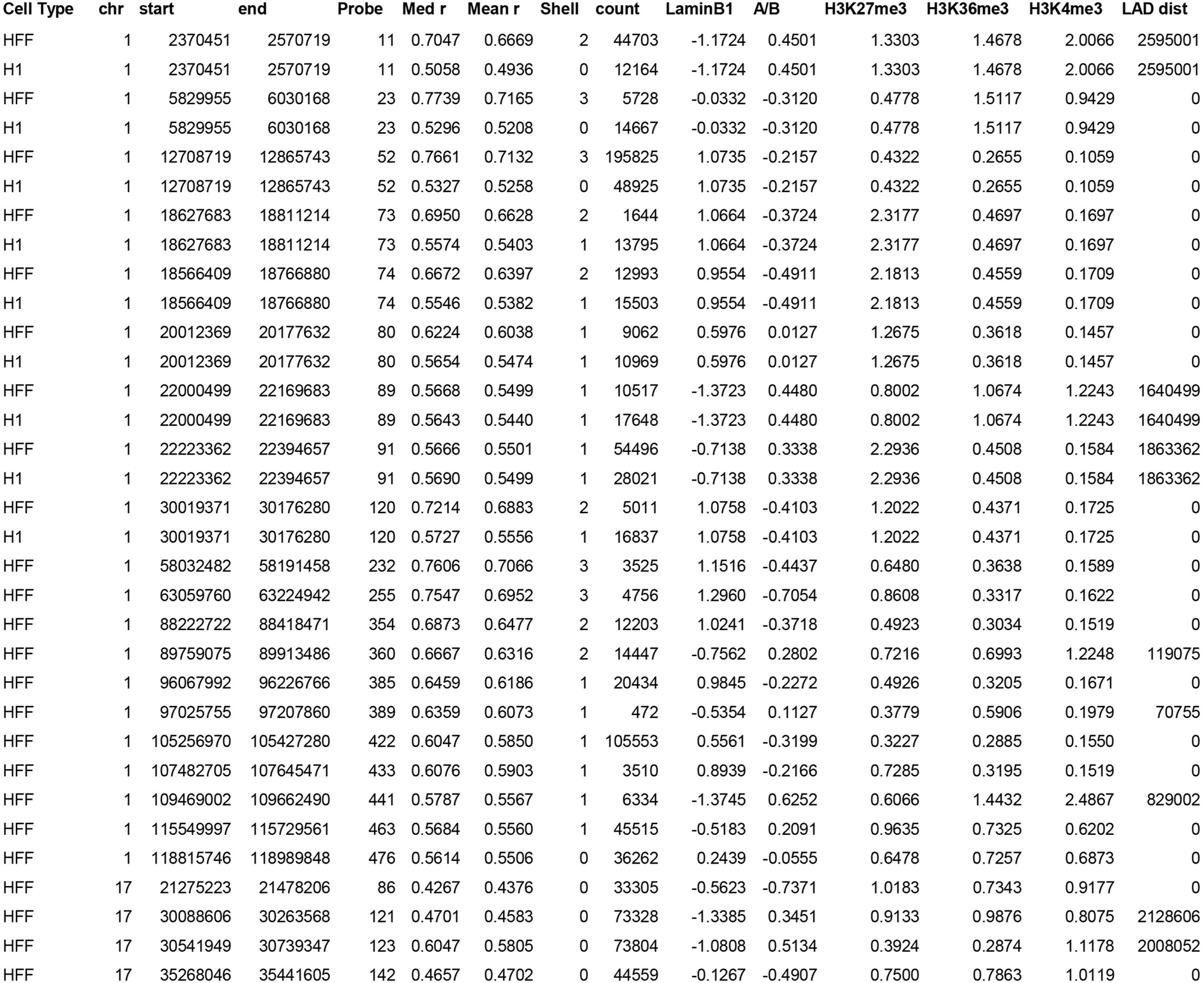
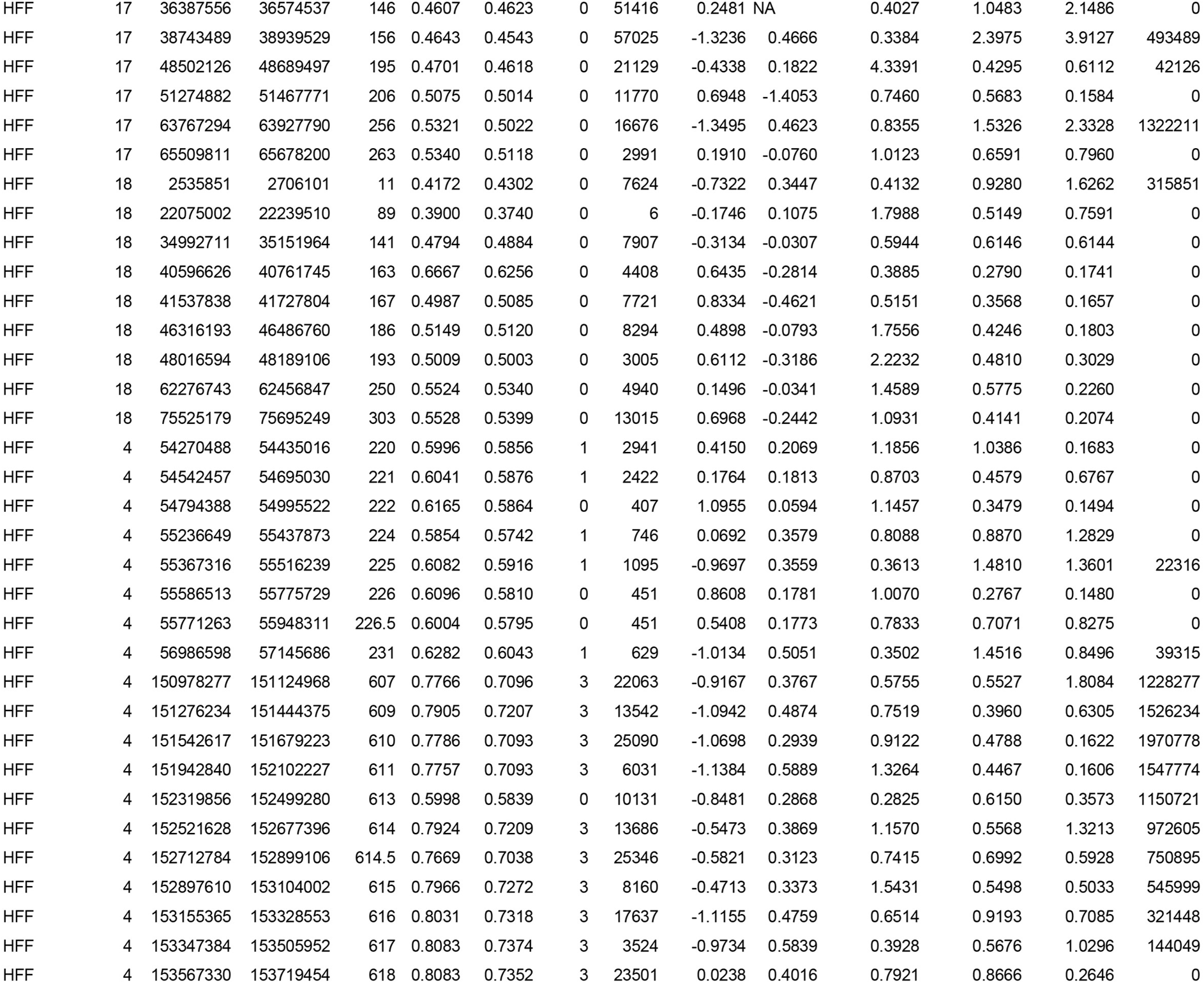
Locus positions and HFF data. Cell Type: Cell type for radial position data. Chr: chromosome. Start: start position of BAC probe. End: end position of BAC probe. Probe: internal probe ID for purposes of figures. Med r: median radial position. Mean r: mean radial position. Shell: most common equi-area radial shell. Count: Number of spots analyzed. LaminB1: Normalized Lamin-B1 Dam-ID signal in HFFs. A/B: Compartment score from micro-C in HFFs. H3K27me3: H3K27me3 ChIP-seq signal in HFFs. H3K36me3: H3K36me3 ChIP-seq signal in HFFs. H3K4me3: HeK4me3 ChIP-seq signal in HFFs. LAD dist: end-to-end distance to nearest LAD in HFFs.

**Table S2:**
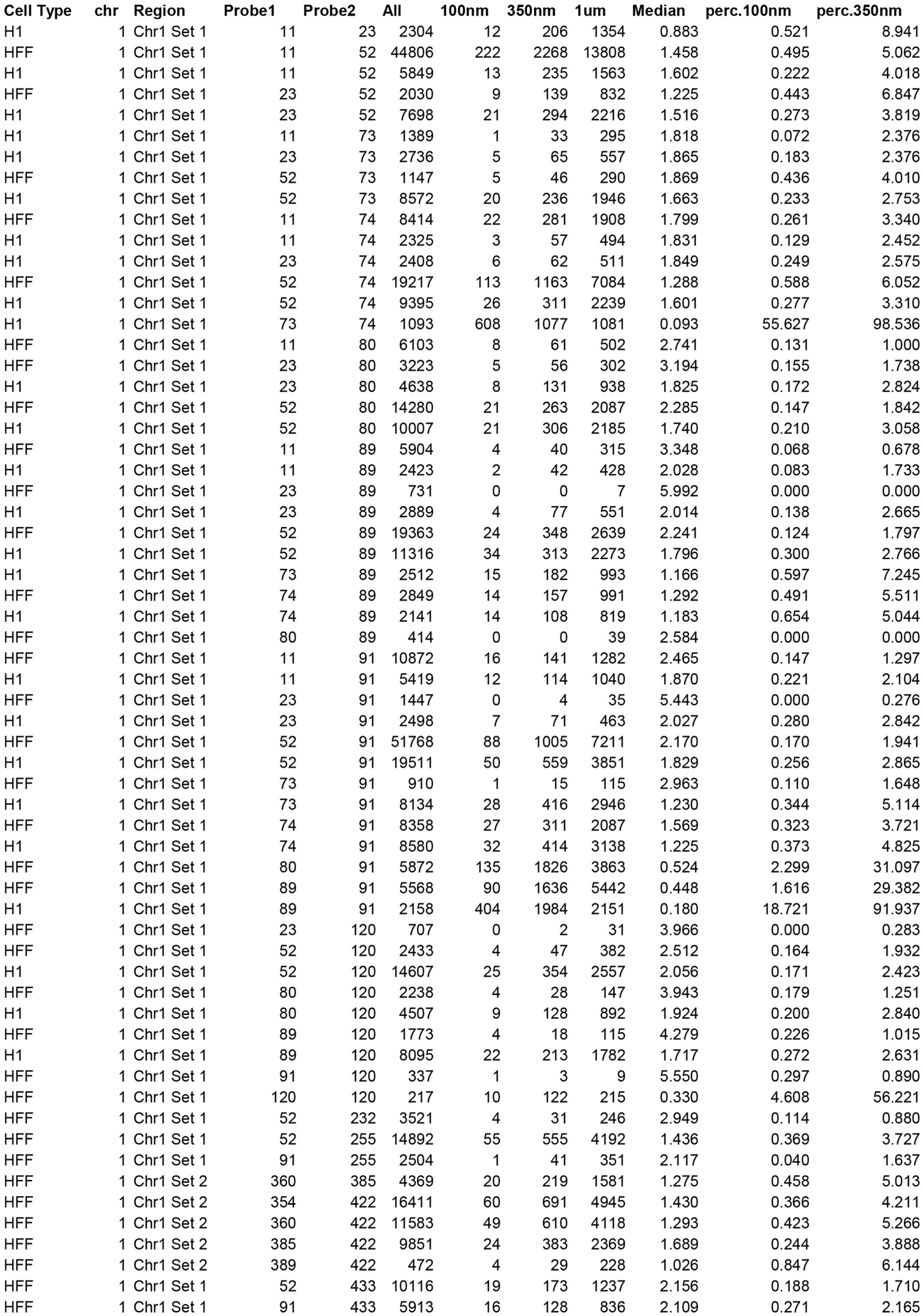
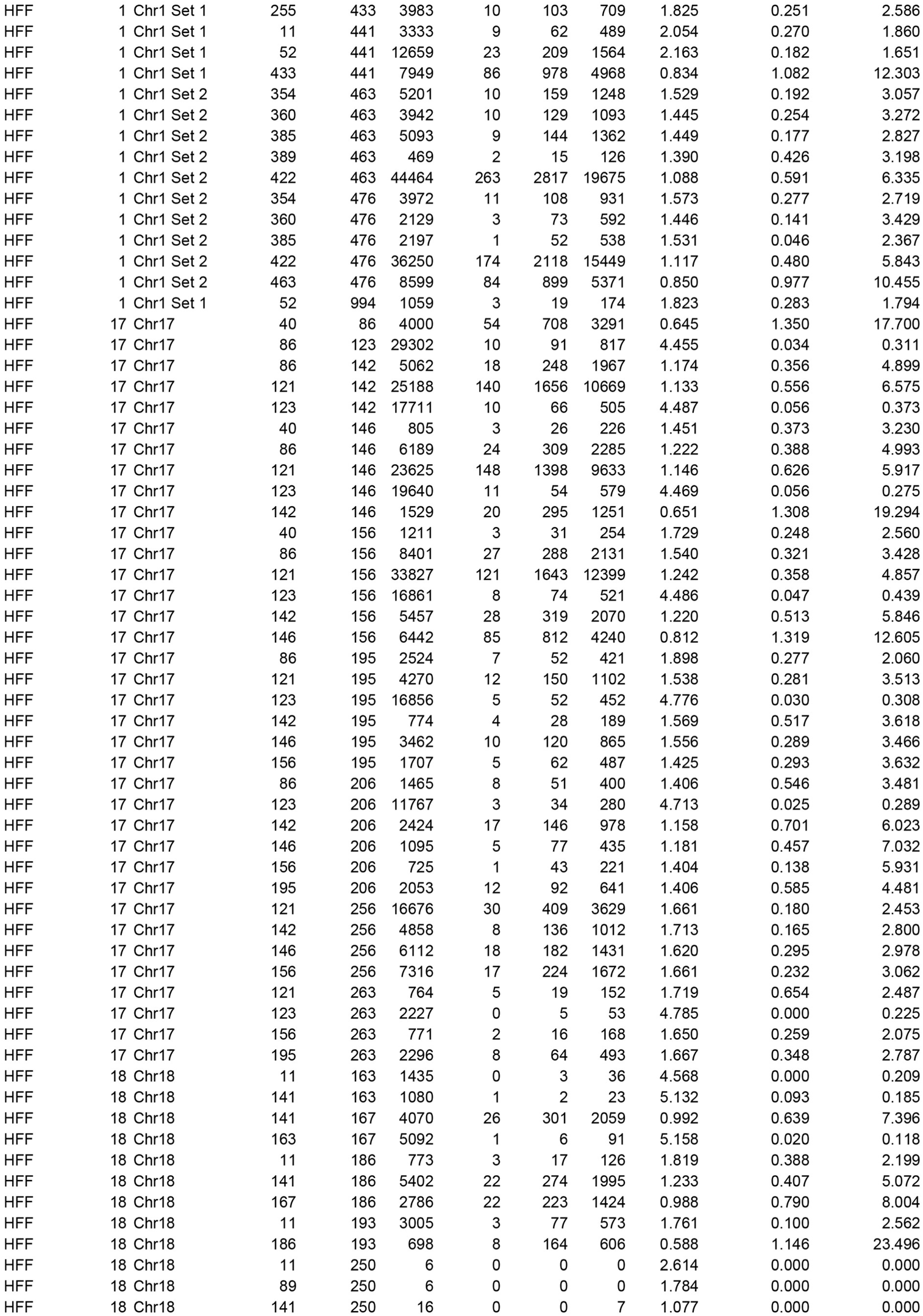
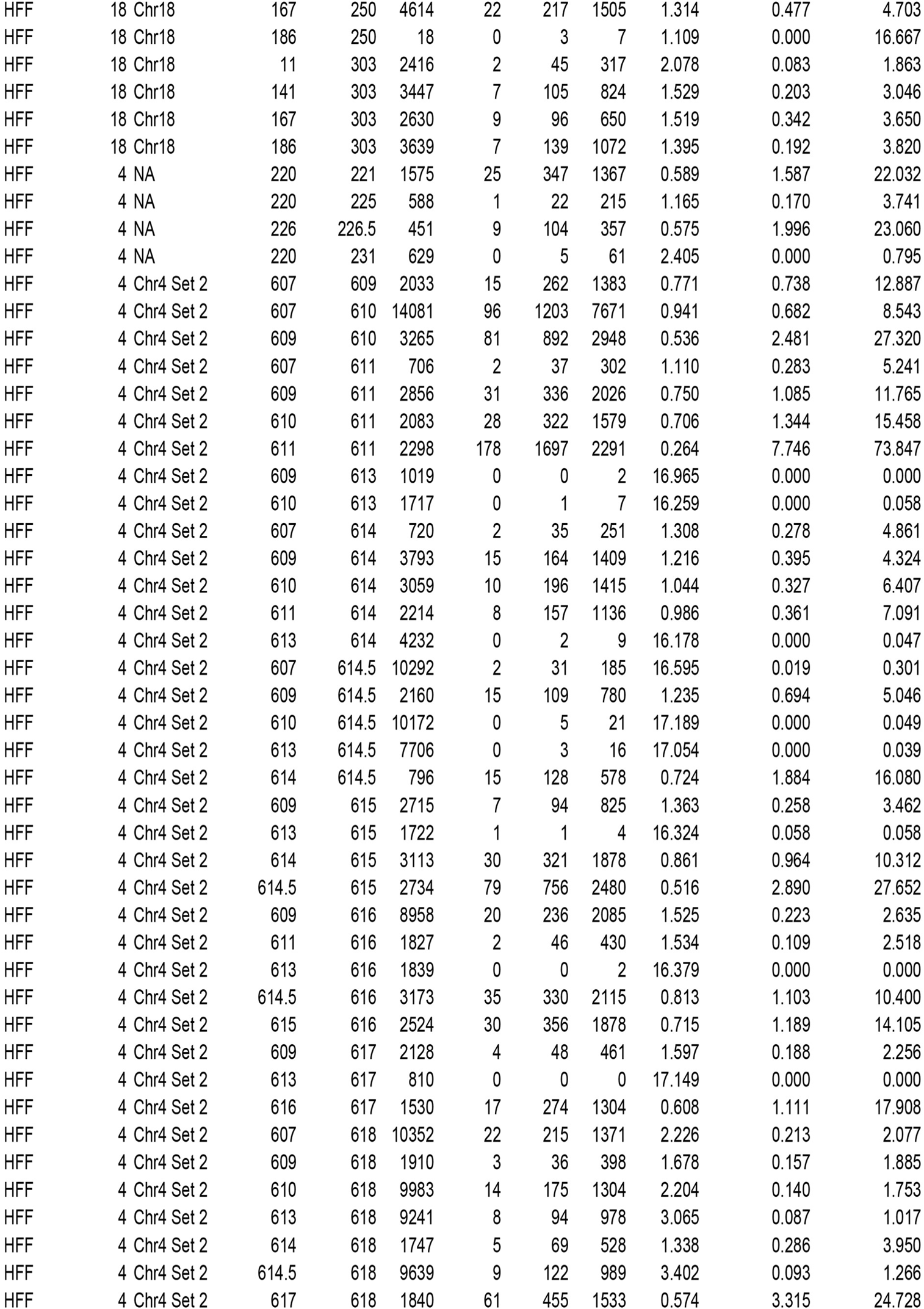
Pairwise distance summary statistics. Cell Type: Cell type. Chr: chromosome. Region: probeset that probe pair belong to (usually chromosome). Probe1: Upstream internal probe ID. Probe2: Downstream internal probe ID. All: Number of spot pairs analyzed. 100nm: Number of spot pairs within 100 nm. 350nm: Number of spot pairs within 350 nm. 1um: Number of spot pairs within 1 micron. Median: Median distance between spots. Perc.100nm: percentage of spot pairs within 100 nm. Perc.350nm: percentage of spot pairs within 350 nm.

